# Half-calcified Calmodulin Promotes Basal Activity and Inactivation of the Calcium Channel Ca_V_1.2

**DOI:** 10.1101/2022.06.24.497440

**Authors:** Peter Bartels, Ian Salveson, Andrea M. Coleman, David E. Anderson, Grace Jeng, Zoila M. Estrada-Tobar, Kwun Nok Mimi Man, Qinhong Yu, Elza Kuzmenkina, Madeline Nieves-Cintron, Manuel F. Navedo, Mary C. Horne, Johannes W. Hell, James B. Ames

**Author notes:** PB, IS, AMC and DEA contributed equally to this work. To whom correspondence should be addressed: (MCH) Department of Pharmacology, University of California at Davis, Davis CA 95616. (JWH) Department of Pharmacology, University of California at Davis, Davis CA 95616. (JBA) Department of Chemistry, One Shields Avenue, University of California, Davis, CA 95616. Author contributions: M.C.H, J.W.H. and J.B.A. designed research and, with input from other authors, wrote the paper; P.B., I.S., A.M.C., D.E.A., G.J., Z.M.E.-T., K.N.M.M., M.N.-C., M.F.N., M.C.H. and J.B.A. performed experiments; P.B., I.S., A.M.C., D.E.A., Q.Y., E.K., M.N.-C., M.F.N., M.C.H, J.W.H., and J.B.A. analyzed data.

## Abstract

The L-type Ca^2+^ channel Ca_V_1.2 controls gene expression, cardiac contraction, and neuronal activity. Calmodulin (CaM) governs Ca_V_1.2 open probability (Po) and Ca^2+^-dependent inactivation (CDI) but the mechanisms remain unclear. We identified a half Ca^2+^-saturated CaM species (Ca_2_/CaM) with Ca^2+^ bound solely at the third and fourth EF-hands (EF3 and EF4) under resting Ca^2+^ concentrations (50-100 nM) that constitutively pre-associates with Ca_V_1.2 to promote Po and CDI. We present an NMR structure of a complex between the Ca_V_1.2 IQ motif (residues 1644-1665) and Ca_2_/CaM_12’_, a calmodulin mutant in which Ca^2+^ binding to EF1 and EF2 is completely disabled. The CaM_12’_ N-lobe does not interact with the IQ motif. The CaM_12’_ C-lobe bound two Ca^2+^ ions and formed close contacts with IQ residues I1654 and Y1657. I1654A and Y1657D mutations impaired CaM binding, CDI, and Po, as did disabling Ca^2+^ binding to EF3 and EF4 in the CaM_34_ mutant when compared to wildtype CaM. Accordingly, a previously unappreciated Ca_2_/CaM species promotes Ca_V_1.2 Po and CDI identifying Ca_2_/CaM as an important mediator of Ca signaling.

## INTRODUCTION

Ca_V_1.2 is the main L-type channel in heart, blood vessels and brain (Ghosh, Syed et al., 2017, Hell, Westenbroek et al., 1993). Ca^2+^ influx through Ca_V_1.2 triggers cardiac contraction, regulates arterial tone (Ghosh et al., 2017), mediates synaptic long-term potentiation (Moosmang, Haider et al., 2005, Qian, Patriarchi et al., 2017), controls neuronal excitability (Berkefeld, Sailer et al., 2006), and mediates Ca^2+^-dependent gene expression (Cohen, Suutari et al., 2018). Defects in inactivation of Ca_V_1.2 causes Timothy syndrome, a rare congenital abnormality leading to lethal arrhythmias, autistic-like behaviors and immune deficiency (Splawski, Timothy et al., 2004). Thus, defining mechanisms of Ca_V_1.2 regulation is highly relevant for understanding its physiological and pathological functions.

Ca^2+^ influx through Ca_V_1.2 triggers a rapid negative feedback mechanism by inducing channel inactivation called Ca^2+^-dependent inactivation (CDI) (Peterson, DeMaria et al., 1999, Zuhlke, Pitt et al., 1999). CDI is mediated by calmodulin (CaM) (Peterson et al., 1999) that is pre-associated with Ca_V_1.2 under basal Ca^2+^ conditions ([Ca^2+^]_i_ = 100 nM) (Erickson, Alseikhan et al., 2001, Erickson, Liang et al., 2003). Ca^2+^-free apoCaM has been suggested to be pre-associated with Ca_V_1.2 (Findeisen, Rumpf et al., 2013) and the closely related Ca_V_1.3 (Ben Johny, Yang et al., 2013). However, under physiological conditions, apoCaM binds to the isolated Ca_V_1.2 IQ-motif with a dissociation constant (K_D_) equal to 10 μM (Evans, Hell et al., 2011, Turner, Anderson et al., 2020) and ~1 μM for full-length Ca_V_1.2 (Erickson et al., 2003). The concentration of free apoCaM is <100 nM in neurons and cardiomyocytes (Turner et al., 2020, Wu & Bers, 2007). Accordingly, the fractional binding of Ca_V_1.2 to apoCaM is predicted to be less than 10% and may not be the prevalent CaM species bound to Ca_V_1.2 or the closely related Ca_V_1.3 under basal conditions as proposed previously (Adams, Ben-Johny et al., 2014, Ben Johny et al., 2013, Findeisen et al., 2013).

To fill a critical gap in our understanding of how CaM governs Ca_V_1.2 function, we used NMR structural analysis, protein biochemistry, and patch-clamp electrophysiology of wild-type and mutated Ca_V_1.2 bound to CaM. Our studies uncovered a half-calcified form of CaM (with Ca^2+^ bound solely at EF3 and EF4, called Ca_2_/CaM) that is functionally pre-associated with Ca_V_1.2 under basal conditions. The NMR structure of Ca_2_/CaM bound to the Ca_V_1.2 IQ-motif (residues 1644-1664) suggests that the Ca^2+^-bound CaM C-lobe (residues F93, M110, L113, M125) forms intermolecular interactions with the side chain atoms from Ca_V_1.2 residues (Y1649, I1654, Y1657 and F1658), whereas the Ca^2+^-free CaM N-lobe does not interact with the IQ motif. Electrophysiology data of key mutants of Ca_V_1.2 (I1654A and Y1657E) contrasted with the earlier findings for the K1662E mutant along with the consequences of ectopic expression of CaM_34_ all suggest that Ca_2_/CaM, rather than apoCaM, pre-associates with Ca_V_1.2 under basal conditions to augment channel open probability (Po) and mediate rapid CDI.

## RESULTS

### A CaM intermediate with two Ca^2+^ bound

ITC studies have suggested that apoCaM binds to the IQ peptide with sub-micromolar affinity in the absence of salt (Findeisen et al., 2013). However, in the presence of physiological salt levels, apoCaM binds to the Ca_V_1.2 IQ-motif with a dissociation constant (K_D_) of 10 micromolar (Evans et al., 2011, Turner et al., 2020). Earlier work suggests that binding of apoCaM to full length Ca_V_1.2 is ~10 times stronger than binding to the IQ segment (Erickson et al., 2003). Collectively, these data suggest that apoCaM binds to full-length Ca_V_1.2 with a K_D_ of ~1 μM, which is outside the physiological concentration range of free CaM (<100 nM) in neurons and cardiomyocytes (Turner et al., 2020, Wu & Bers, 2007), implying low fractional binding. Furthermore, the recent NMR structure of apoCaM bound to the Ca_V_1.2 IQ-motif revealed an intermolecular salt bridge involving Ca_V_1.2 residue K1662, and the K1662E mutation significantly and selectively weakened apoCaM binding to Ca_V_1.2 (Turner et al., 2020). At the same time, the K1662E mutation does not affect single-channel Po (Turner et al., 2020). These previous results suggest that apoCaM may not be the main CaM species to support Ca_V_1.2 activity under basal conditions as proposed previously (Adams et al., 2014, Ben Johny et al., 2013, Findeisen et al., 2013). The current study tested the hypothesis that the Ca_V_1.2 channel may pre-associate mostly with a CaM species that is half saturated with Ca^2+^ under basal Ca^2+^ conditions ([Ca^2+^]_i_ = 100 nM).

In support of our hypothesis, we find that that IQ binding to CaM causes a more than 10-fold increase in the apparent Ca^2+^ affinity, which allows Ca^2+^ to bind to the CaM C-lobe under basal conditions (*Suppl. Fig. 1*). On the basis of previous binding data (Ames, 2021, Evans et al., 2011, Gilli, Lafitte et al., 1998), the C-lobe under basal conditions is predicted to bind two Ca^2+^ to form a half-calcified state (called Ca_2_/CaM) in which the N-lobe is devoid of Ca^2+^ (Ames, 2021). Indeed, the C-lobe binds Ca^2+^ as well as the IQ motif with 10-fold higher affinity than the N-lobe (Evans et al., 2011, Gilli et al., 1998). Using the binding constants from (Evans et al., 2011, Gilli et al., 1998), the relative concentrations of apoCaM, CaM intermediate (Ca_2_/CaM), and Ca^2+^-saturated CaM (Ca_4_/CaM) each bound to the IQ as a function of free Ca^2+^ concentration are shown in *Suppl. Fig. 1*. The Ca_2_/CaM intermediate species (red trace in *Suppl. Fig. 1A* has a significant occupancy of ~50% at 100 nM Ca^2+^ concentration (basal Ca^2+^ level). Since the apoCaM N-lobe (CaMN) does not bind to IQ under physiological conditions (Evans et al., 2011), IQ must instead be bound to the C-lobe (CaMC) of Ca_2_/CaM. Using binding constants from (Evans et al., 2011, Gilli et al., 1998)) we calculate that CaMC-IQ (Scheme 1) and CaMN-IQ (Scheme 2) have apparent K_D_ values for Ca^2+^-binding of 100 nM and 1.0 μM, respectively:

#### Scheme 1

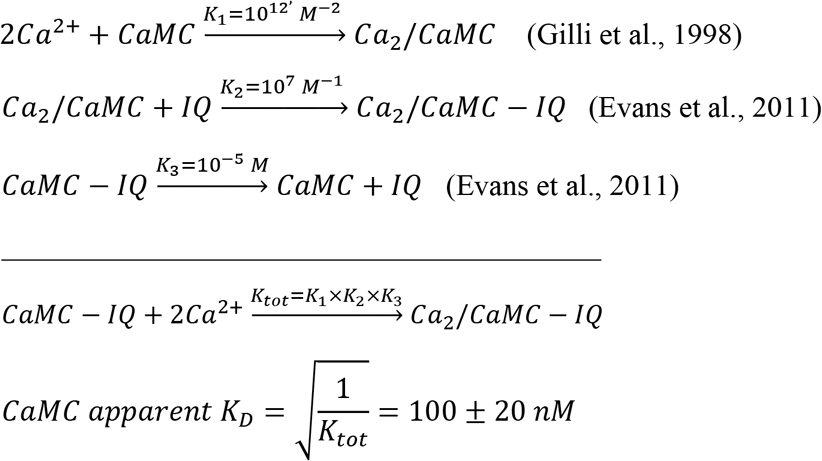

#### Scheme 2

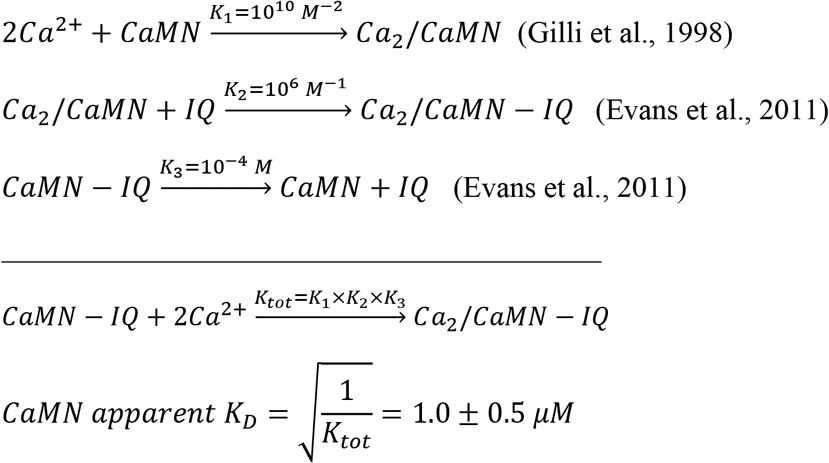

Thus, the CaM C-lobe is calculated to have a 10-fold higher apparent Ca^2+^ affinity compared to CaM N-lobe. This calculation implies that ~50% of CaM-IQ complex will have Ca^2+^ bound to its C-lobe under basal conditions ([Ca^2+^]_i_ = 100 nM), whereas the N-lobe should be devoid of Ca^2+^. To test this prediction, we prepared a CaM mutant (D21A/D23A/D25A/E32Q/D57A/D59A/N61A/E68Q, called CaM_12’_) that completely disabled Ca^2+^ binding to EF1 and EF2, but retained normal Ca^2+^ binding to EF3 and EF4. The apparent Ca^2+^ affinity of CaM_12’_ in the presence of saturating IQ peptide under physiological conditions was measured by isothermal titration calorimetry (ITC) (*Suppl*. Fig. 2). The ITC isotherm is biphasic, suggesting possible sample heterogeneity. The major binding component (N_2_ = 1.7 ±0.3 Ca^2+^/protein; *Suppl. Tab.1*) represents binding of two Ca^2+^ to CaM_12’_-IQ as defined by K_2_, ΔH_2_ and N_2_ (*Suppl. Tab.1*). The other isotherm component is non-stoichiometric (N1 = 0.2 ±0.1 Ca^2+^/protein) and may be an artifact of IQ dimerization or other sample heterogeneity. Fitting the ITC isotherm with a 2-site model reveals a Ca^2+^-binding apparent K_D_ 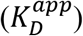 of 60 ±20 nM (*Suppl. Tab.1*), which agrees within experimental error with the predicted value in Scheme 1 and with previously measured values of 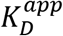 obtained by UV fluorescence (Halling, Georgiou et al., 2009). The relatively high apparent Ca^2+^ affinity 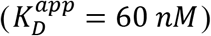 implies that at least 50% of the CaM/IQ complex will have Ca^2+^ bound to EF3 and EF4 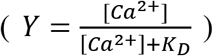 at basal Ca^2+^concentrations (~100 nM). This analysis predicts that slightly more than half of the Ca_V_1.2 channels should be pre-associated with the CaM intermediate, Ca_2_/CaM under basal conditions.

### Half-calcified CaM represented by CaM_12’_

The concentration profiles in *Suppl*. Fig. 1A show that half saturated CaM (Ca_2_/CaM) co-exists in an equilibrium mixture with apoCaM and Ca^2+^-saturated CaM (Ca_4_/CaM). Therefore, under basal conditions, Ca_2_/CaM cannot be resolved from the other CaM species. To isolate the half Ca^2+^ saturated species, we performed structural studies on the CaM mutant (D21A/D23A/D25A/E32Q/D57A/D59A/N61A/E68Q, called CaM_12’_) that completely disables Ca^2+^ binding to EF1 and EF2, but retains Ca^2+^ binding to EF3 and EF4. The NMR assignments of Ca^2+^-bound CaM_12’_ bound to the IQ peptide (Ca_2_/CaM_12’_-IQ) reveal two downfield NMR peaks assigned to G99 (EF3) and G135 (EF4) that indicate Ca^2+^ is bound to EF3 and EF4 (Salveson, Anderson et al., 2019). The corresponding Gly residues in EF1 (G26) and EF2 (G62) do not exhibit downfield amide resonances, indicating that EF1 and EF2 in Ca_2_/CaM_12’_-IQ are both devoid of Ca^2+^.

The NMR spectrum of Ca_2_/CaM_12’_-IQ is a hybrid of the spectra of Ca^2+^-bound and Ca^2+^-free CaM_12’_ (Figs. 1A-B). The chemical shifts assigned to the CaM_12’_ C-lobe (residues 80-149) of Ca_2_/CaM_12’_-IQ (peaks labeled red in Fig. 1A) are nearly identical to those of the isolated Ca^2+^-bound CaM C-lobe bound to IQ (blue peaks in Fig. 1A). NMR peaks assigned to CaM_12’_ N-lobe (residues 1-79) of Ca_2_/CaM_12’_-IQ are similar to those of apoCaM in the absence of IQ (black peaks in Fig. 1B), indicating that the CaM_12’_ N-lobe is Ca^2+^-free and does not interact with the IQ peptide. Thus, only the C-lobe but not N-lobe residues in Ca_2_/CaM_12’_ exhibit IQ-induced spectral shifts.

**Figure 1.**
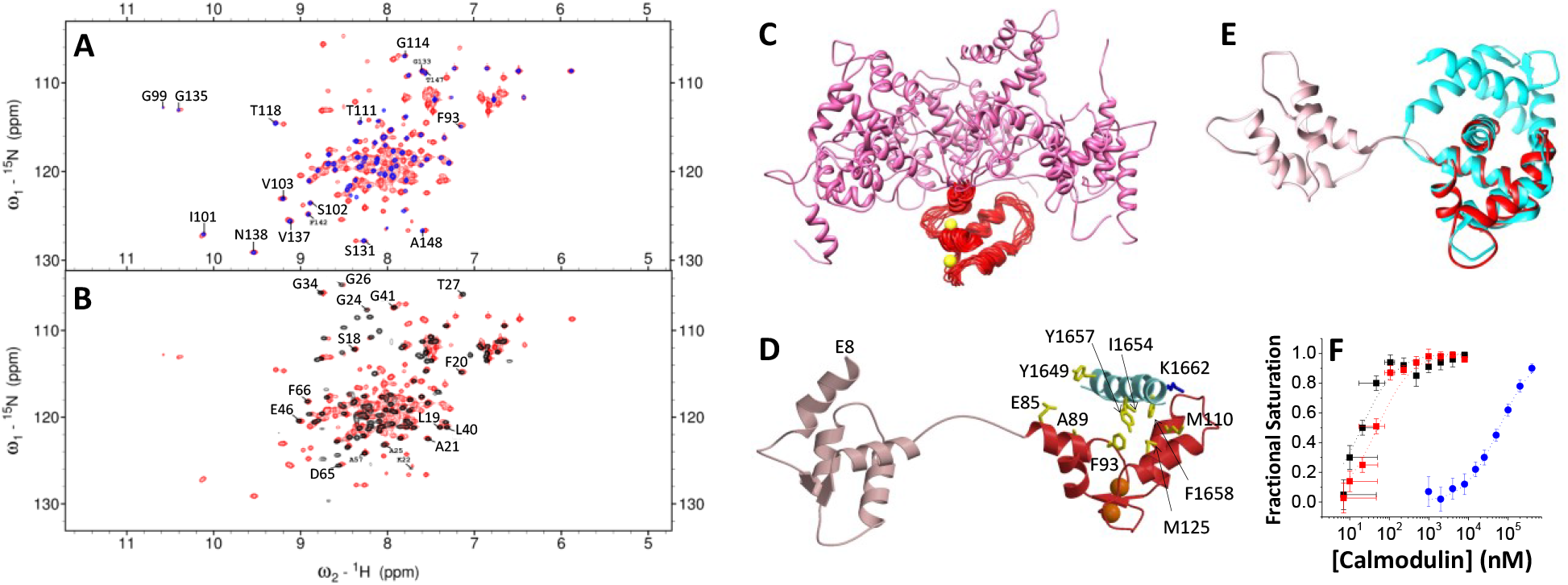
NMR-derived structures of Ca_2_/CaM_12’_-IQ. (**A**) ^15^N-^1^H HSQC NMR spectrum of ^15^N-labeled Ca_2_/CaM_12’_ bound to unlabeled IQ (red) is overlaid with the spectrum of Ca^2+^-bound CaM^WT^ C-lobe/IQ complex (blue). (**B**) NMR spectrum of ^15^N-labeled Ca_2_/CaM_12’_ bound to unlabeled IQ (black) is overlaid with the spectrum of Ca^2+^-free CaM_12’_ (red). (**C**) Ensemble of 10 lowest energy NMR structures of Ca_2_/CaM_12’_ (PDB ID: 7L8V). Main chain structures are depicted by a ribbon diagram. Structures of the C-lobe (residues 85-149) are overlaid and highlighted in red; N-lobe structures (residues 1-84) are highlighted in pink. Bound Ca^2+^ ions are yellow. Structural statistics are given in Table 1. (**D**) The lowest energy structure of Ca_2_/CaM_12’_-IQ complex is shown as a ribbon diagram of Ca_2_/CaM_12’_ bound to the IQ peptide (cyan). The CaM N-lobe and C-lobe are highlighted pink and red, respectively. Side-chain atoms of key residues are depicted by sticks and are colored yellow and blue. (E) Overlay of the NMR structure of Ca_2_/CaM_12’_-IQ (C-lobe in red) with the crystal structure of Ca_4_/CaM (cyan, 2BE6). The C-lobe structures overlay with an RMSD of 1.8 Å. (**F**) Fluorescence polarization assay showing the binding of half Ca^2+^-saturated CaM mutant (Ca_2_/CaM_12’_) with fluorescently labeled IQ peptides (wildtype: black; K1662E: red; both: K_D_ < 100 nM), and of apoCaM binding to Y1657D (blue, K_D_ = 60 μM).

### NMR structure of Ca_2_/CaM_12’_-IQ

NMR spectral assignments for Ca_2_/CaM_12’_-IQ were reported previously (BMRB accession number 27692) (Salveson et al., 2019). These previous NMR assignments were used in the current study to obtain NMR-derived structural restraints from NOESY and residual dipolar coupling (RDC) data (*Suppl*. Fig. 3). NMR structures of Ca_2_/CaM_12’_-IQ were then calculated on the basis of distance restraints derived from analysis of NOESY (Clore & Gronenborn, 1998) and long-range orientational restraints derived from RDC data (Tjandra & Bax, 1997) as described in the Experimental Procedures. The final NMR-derived structures of Ca_2_/CaM_12’_ are overlaid in Fig. 1C and structural statistics summarized in Table 1. The two domains of Ca_2_/CaM_12’_ (N-lobe in pink and C-lobe in red, Fig. 1C) are separately folded and non-interacting as was seen previously for the NMR structures of apoCaM (Finn, Evenas et al., 1995, Kuboniwa, Tjandra et al., 1995, Zhang, Tanaka et al., 1995). The overall precision of the NMR ensemble is expressed by a root-mean-square deviation (RMSD) of 0.83±0.09 Å calculated from the coordinates of the main chain atoms in the C-lobe (Fig. 1C), and 0.9±0.1 Å from the main chain atoms in the N-lobe. The lowest energy NMR structure of Ca_2_/CaM_12’_ bound to the IQ peptide is shown in Fig. 1D. The quality of the NMR structures of Ca_2_/CaM_12’_-IQ was assessed using PROCHECK-NMR (Laskowski, Rullmann et al., 1996), which shows that 93% of the residues occur in the allowed or favorable regions from the Ramachandran plot. The NMR structure of the Ca^2+^-bound CaM C-lobe (residues 80-149) of Ca_2_/CaM_12’_-IQ (dark red in Figs. 1D and 1E) looks similar to that observed in the crystal structure of Ca^2+^-saturated CaM bound to the IQ (cyan in Fig. 1E) (Van Petegem, Chatelain et al., 2005). The structure of the Ca^2+^-free CaM N-lobe (residues 1-78) of Ca_2_/CaM_12’_-IQ (light red in Fig. 1D) adopts a closed conformation and looks similar to that of apoCaM (Zhang et al., 1995). The IQ peptide was verified by NMR to have a helical conformation (cyan in Fig. 1D). In the Ca_2_/CaM_12’_-IQ structure (Fig. 1D), the IQ residues (Y1649, I1654, Y1657 and F1658) point toward CaM and make extensive contacts with CaM C-lobe residues (E85, A89, F93, M110, L113, M125). The IQ peptide in the Ca_2_/CaM_12’_-IQ structure does not make any contacts with the Ca^2+^-free N-lobe, in contrast to the crystal structure of Ca_4_/CaM_12’_-IQ (Fallon, Baker et al., 2009, Fallon, Halling et al., 2005, Van Petegem et al., 2005) where IQ aromatic residues (F1648, Y1649 and F1652) make extensive contacts with N-lobe residues (F13, F69, M73).

**Table 1.**
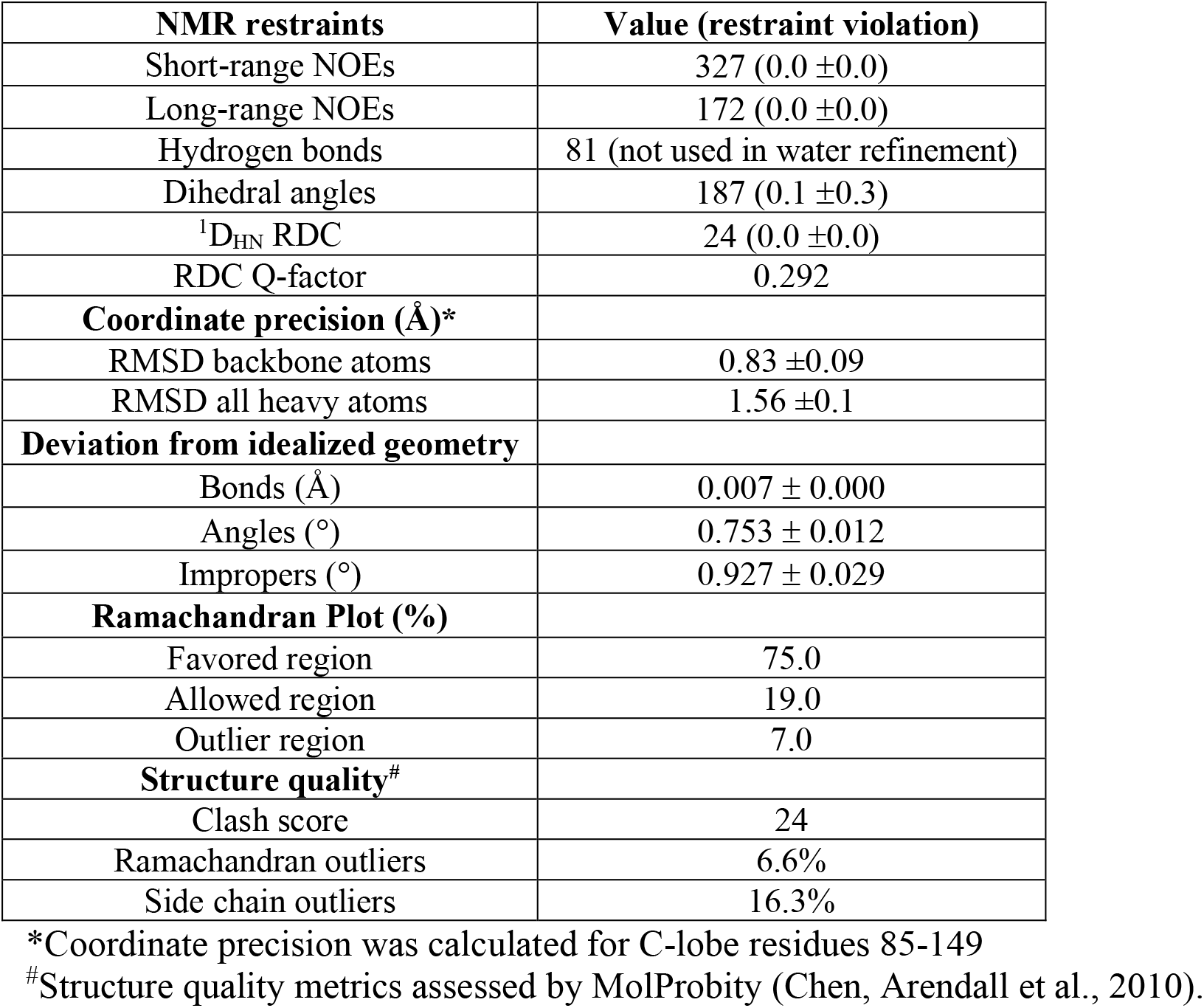
NMR Structural Statistics for Ca_2_/CaM_12’_-IQ

### IQ residue K1662 interacts with apoCaM more strongly than Ca_2_/CaM_12’_

The NMR structure of Ca_2_/CaM_12’_-IQ (Fig. 1D) looks quite different from the recent NMR structure of apoCaM bound to IQ (Turner et al., 2020). In the apoCaM-IQ structure, K1662 forms intermolecular salt bridges with CaM residues, E85 and E88. By contrast, K1662 is mostly solvent exposed in the Ca_2_/CaM_12’_-IQ structure and does not contact either E85 or E88 (Fig. 1D). This analysis predicts that the Ca_V_1.2 mutation K1662E weakens binding to apoCaM more than it does to Ca_2_/CaM_12’_. Because the K1662E peptide (IQ^K1662E^) was not soluble enough for ITC with Ca_2_/CaM_12’_, we used fluorescence polarization (FP) to measure binding affinity in the nanomolar range. As predicted, titration of the IQ peptides with Ca_2_/CaM_12’_ reached full saturation at 100 nM Ca_2_/CaM_12’_ indicating a K_D_ < 100 nM for both, IQ^WT^ and IQ^K1662E^ (Fig. 1F). It was not possible to more accurately determine the actual K_D_ because the IQ peptide concentration in Fig. 1F had to be 100 nM due to limited detection sensitivity. This concentration is much larger than the K_D_ for IQ^WT^ (16 nM in Suppl. Table 2) and apparently also for IQ^K1662E^ as binding was clearly saturated at 100 nM for both peptides. The free concentrations of Ca_2_/CaM_12’_ ([*Ca*_2_/*CaM*_12′_]*_free_* = [*Ca*_2_/CaM_12′_]*_total_* – [*IQ*] × (*fractional saturation*)) are within the sample noise level during the first half of the titration when [*Ca*_2_/*CaM*_12′_]*_free_* < 100 *nM* (see SD bars in Fig. 1F). During the second half of the titration, [*Ca*_2_/*CaM*_12′_]*_free_* was above the noise level and the titration curves show clear saturation at 100 nM providing an upper limit of 100 nM for the K_D_ of both, IQ^WT^ and IQ^K1662E^, consistent with the 16 nM K_D_ for IQ^WT^ as seen by ITC (Suppl. Table 2). As a result, Ca_2_/CaM_12’_ can bind to IQ^K1662E^ in the nanomolar range in contrast to apoCaM, which binds to IQ^K1662E^ with a K_D_ in the high micromolar range (60 μM) that is 6-fold higher than that of IQ^WT^ (Turner et al., 2020). Thus, the K1662E mutation weakens IQ binding to apoCaM to a degree that is outside the physiological range of its concentration (Wu & Bers, 2007) (<100 nM), in contrast to the nanomolar binding of IQ^K1662E^ with Ca_2_/CaM_12’_ (Fig. 1F). Accordingly, the K1662E mutation can be used to selectively disable apoCaM binding to Ca_V_1.2, while retaining Ca_V_1.2 binding to Ca_2_/CaM.

### IQ residues Y1649,I1654, Y1657 and F1658 interact with Ca_2_/CaM_12’_

The NMR structure of Ca_2_/CaM_12’_-IQ reveals intermolecular contacts with IQ residues, Y1649, I1654, Y1657 and F1658 that are each located on the same side of the IQ helix pointing toward the Ca^2+^-occupied C-lobe of Ca_2_/CaM_12’_ (Fig. 1D). As predicted by this analysis, the IQ peptide mutants IQ^Y1649A^, IQ^F1654A^, IQ^Y1657D^, IQ^F1658D^ each exhibited weaker binding to Ca_2_/CaM_12’_ compared to IQ^WT^. The K_D_ was 16 ±5 nM for IQ^WT^, 26 ±5 nM for IQ^Y1649A^, 60 ±10 nM for IQ^I1654A^, 8000 ±10 nM for IQ^Y1657D^, 4000 ±10 nM for IQ^F1658D^, and 32 ±5 nM for IQ^F1658A^ (*Suppl*. Figs. 2B-F and Table S2). These findings validate our structural analysis and verify that Y1657 makes the strongest contact with CaM.

**Figure 2.**
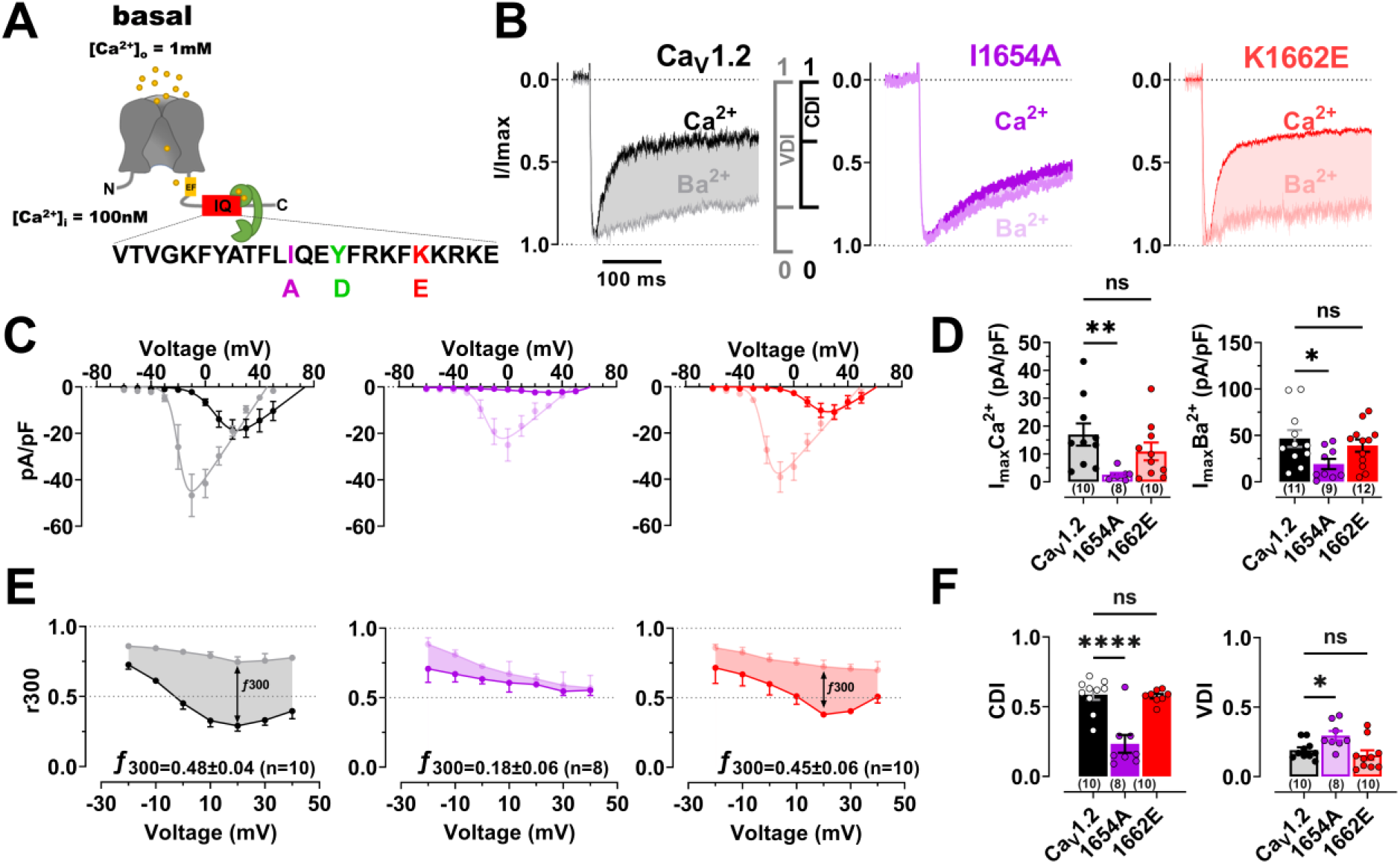
Effects of IQ mutants I1654A and K1662E on Ca_V_1.2 activity and inactivation. (**A**) Topology of the hypothetical Ca_V_1.2 Ca^2+^ channel pore and localization of the IQ domain and its mutations in the α_1_1.2 subunit. At rest with [Ca^2+^]_i_ ≤100 nM the C-lobe (green) of half-calcified Ca^2+^/CaM is predicted to bind to the C-terminal portion of the IQ motif, making hydrophobic contacts with I1654 and Y1657 but not with K1662. (**B-F**) HEK 293T/17 cells were transfected with α_1_1.2, α_2_δ1, and β_2A_. Shown are representative whole-cell current traces (**B**), population data of current-voltage relationships (I/V curves) (**C**), their respective peak current density plots (**D**), and currents remaining after 300 ms of depolarization (r300; bottom) of I_Ba_ (10 mM Ba^2+^; grey or light colors) and I_Ca_ (10 mM Ca^2+^; black or dark colors), for WT (black), I1654A (purple) and K1662E (red) (**E**). Peak currents in B were normalized to the respective current maxima (Imax). Shaded areas indicate differences between I_Ba_ and I_Ca_ as read out for CDI (f300: difference between I_Ba_ and I_Ca_ remaining after 300 ms). Quantification of peak current densities of I_Ba_ and I_Ca_ at potential of respective Imax reveals a strong decrease in current density for I1654A but not K1662E versus WT (**D**). (**F**) Quantification of CDI and VDI reveals a strong decrease in CDI for I1654A but not K1662E versus WT. Mutations show modest, statistically non-significant effects on VDI versus WT. Numbers in parenthesis under bars reflect n independent recordings and error bars SEM (*p<0.05, p**<0.01, and ****p<0.0001, One-Way ANOVA with Bonferroni correction).

### The K1662E mutation affects binding of apoCaM but not CDI of Ca_V_1.2

The above analysis suggests that K1662E retains binding to Ca_2_/CaM_12’_ but not apoCaM under physiological conditions (i.e., with free CaM < 100 nM (Wu & Bers, 2007)) (Fig. 1F). This differential effect informs interpretation of recently published data that showed that the K1662E mutation has no effect on Po (Turner et al., 2020), while the I1654A mutation, which affects binding of both apoCaM and Ca/CaM, decreased Po by 6-fold (Turner et al., 2020). A similar effect has been seen for an analogous Ile to Ala mutation in the closely related Ca_V_1.3 (Adams et al., 2014). Collectively these findings suggest that CaM promotes Po when it forms a complex with Ca_V_1.2 with Ca^2+^ bound to EF3 and EF4 to give rise to a half-saturated Ca_2_/CaM state in this complex. To further test the idea of pre-association of half Ca^2+^-saturated Ca_2_/CaM with Ca_V_1.2 at basal Ca^2+^ concentrations, we wanted to compare CDI of Ca_V_1.2^K1662E^ with WT and also Ca_V_1.2^I1654A^, which served as a well-established reference point for loss of CDI (Adams et al., 2014, Ben Johny et al., 2013, Peterson et al., 1999). For that purpose, we measured whole-cell current density for I_Ba_ and I_Ca_. Consistent with the earlier Po analysis, I_Ba_ and I_Ca_ were reduced by the I1654A but not K1662E mutation (Fig. 2A-D; *Suppl*. Table 3A). Strikingly, the K1662E mutation had no significant effect on CDI (nor on VDI), in contrast to the I1654A mutation, which reduced CDI by ~75% (Fig. 2B,E,F; *Suppl*. Table 3B). The small, remaining CDI seen for the I1654A mutant channel may be due to N-lobe effects such as its binding to the N-terminus of the Ca_V_1.2 α_1_ subunit channel (Dick, Tadross et al., 2008). The differential effect on I_Ba_, I_Ca_, and CDI by the K1662E versus I1654A mutation is consistent with the differential effect of the K1662E versus I1654A mutation on Po (Turner et al., 2020) and suggests that formation of a complex of Ca_V_1.2 with half Ca^2+^-saturated Ca_2_/CaM is important for Po and for predisposing Ca_V_1.2 to CDI.

### The Y1657D mutation strongly affects binding of half-saturated Ca_2_/CaM as well as I_Ba_, I_Ca_, Po, and CDI of Ca_V_1.2

Our new Ca_2_/CaM_12’_-IQ structure indicates that Y1657 makes the most and closest contacts among all IQ residues with Ca_2_/CaM_12’_ (Fig. 1). In support of its central role in mediating this interaction, binding studies indicate that the Y1657D mutation has the strongest negative effect on the affinity of the Ca_2_/CaM_12’_-IQ interaction of all tested IQ peptides (K_D_ for IQ^WT^ is 16 nM and for IQ^Y1657D^ 8 μM; *Suppl*. Tab. 2). The Y1657D mutation decreased whole-cell currents, I_Ba_ and I_Ca_ as well as CDI with no effect on VDI (Fig. 3A-E). Single-channel recordings show a remarkably strong decrease in Po for Y1657D versus WT Ca_V_1.2 (Fig. 3F,G). This loss in Po and CDI is comparable to similarly strong effects for the I1654A mutation on Po (Turner et al., 2020) and CDI (Zuhlke et al., 1999) but the K1662E mutation, which specifically affects apoCaM but not Ca/CaM binding, did not affect Po (Turner et al., 2020) or CDI (Fig. 2). The decrease in Po is also well reflected when calculating the ensemble averages of unitary single-channel currents (Fig. 3F; *Suppl*. Tab. 4). To test whether there is also a change in channel surface expression in addition to a decrease in Po of individual channels, we conducted surface biotinylation experiments. We determined that Ca_V_1.2 surface expression was reduced by almost 50% (Fig. 3H,I), which can explain some, but not all, of the 80% loss in Po.

**Figure 3.**
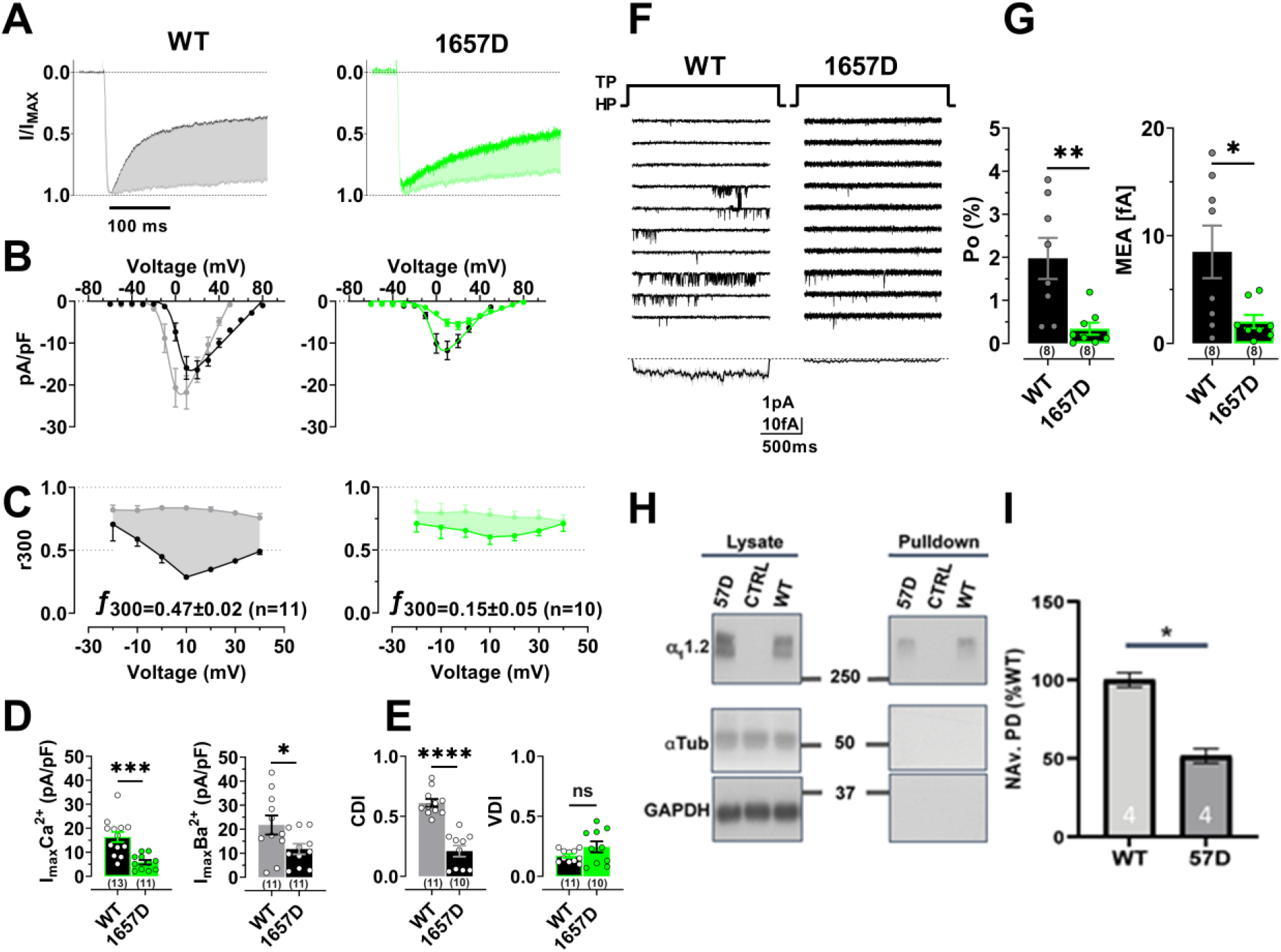
Effects of IQ mutant Y1657D on Ca_V_1.2 activity, inactivation, and surface expression. HEK 293T/17 cells were transfected with α_1_1.2, α_2_δ1, and β_2A_. Shown are representative whole-cell current traces (**A**), population data of I/V curves (**B**) currents remaining after 300 ms of depolarization (r300; bottom) of I_Ba_ (grey or light green) and I_Ca_ (black or dark green), for WT (black) and Y1657D (green) (**C**), and peak current density plots (**D**). Peak currents in A were normalized to the respective current maxima (Imax). Shaded areas indicate differences between I_Ba_ and I_Ca_ as read out for CDI (f300: difference between I_Ba_ and I_Ca_ remaining after 300 ms). (**D**) Quantification of peak I_Ba_ and I_Ca_ at potential of respective Imax reveals a strong decrease in current density for Y1657D versus WT. (**E**) Quantification of CDI and VDI reveals a strong decrease in CDI but not VDI for Y1657D versus WT. (**F**) 10 consecutive representative single-channel traces of WT and Y1657D. Below: mean ensemble average currents (MEA) calculated from a total of 857 superimposed traces for WT (n=8 cells) and 1366 traces for Y1657D (n= 8 cells). (**G**) Quantification of single-channel open probability Po (left) and MEA (right) reveals a strong decrease in channel activity for Y1657D versus WT. (**H**) Surface biotinylation of Ca_V_1.2 was followed by Neutravidin pulldowns and immunoblotting (right panels) with antibodies against the proteins indicated at the left. Left panels show immunoblots of total lysate. Tubulin (α-Tub), and GAPDH were used as loading controls for lysate samples (left) and assessment of membrane integrity (right; all left and right panels were from same gels and exposures). Absence of tubulin and GAPDH immunoreactivity indicates that the biotin reagent did not leak into cells ruling out biotinylation of intracellular proteins. (**I**) Quantification of α_1_1.2 immunosignals in Neutravidin pulldowns (NAv.PD) normalized to WT α_1_1.2 (set to 100%). The 1657D α_1_1.2 mutant exhibits a decrease in surface biotinylation relative to the WT subunit (n = 4; p = 0.0132, two-tailed t-test). Numbers in parenthesis under bars or inside bars reflect “n” independent recordings or pull downs and error bars SEM (*p<0.05, **p<0.01, ***p<0.001 and ****p<0.001, unpaired T-test).

### CaM Intermediate (Ca_2_/CaM) Increases Po of Ca_V_1.2

To further analyze the role of CaM in Po we ectopically expressed CaM in 293T cells. Although this approach has been used before to define the role of CaM in CDI, the level to which exogenous CaM was expressed in these CDI studies had not been thoroughly assessed (Iacobucci & Popescu, 2019). Thus, we investigated whether the expression of CaM_34_ (described by (Peterson et al., 1999)) was sufficient to allow detection of an effect (i.e., many fold greater than endogenous CaM) by immunoblotting extracts of 293T cells transfected with Ca_V_1.2 expression constructs ± WT CaM or CaM_34_ plasmids (*Suppl. Fig. 4*). We found that overexpression of WT compared to endogenous CaM is about ~10 fold, while CaM_34_ is ~20 fold (*Suppl. Fig. 4 A-D*). To test whether ectopic expression of CaM affects levels of endogenous CaM, we expressed YFP-tagged WT CaM or CaM_34_, which migrate at an M_R_ of ~ 45 kDa (verified by anti-YFP immunoblotting; *Suppl. Fig 4 E*). Probing immunoblots with anti-CaM identifies a prominent 45 kDa band and a weaker signal for the endogenous 17 kDa band. Comparison of the 17 kDa band in mock-transfected (no CaM vectors) cell lysate to the same MR immunoreactive band in the CaM plasmid transfected samples did not indicate a significant effect of ectopic CaM on endogenous CaM levels (*Suppl. Fig. 4E,F*).

Consistent with earlier work on Ca_V_1.3 by Adams et al. (Adams et al., 2014), we find that overexpression of WT CaM strongly increases Po by ~300% as compared to expression of Ca_V_1.2 alone (Fig. 4A,B; *Suppl. Tab. 5*). This effect could be due to increased binding of apoCaM, half Ca^2+^-saturated Ca_2_/CaM, or both. Because earlier work did not differentiate between these possibilities (Adams et al., 2014), we tested the effect of ectopic expression of CaM_34_ and found no increase at all in Po as compared to expression of Ca_V_1.2 alone. This result demonstrates that Ca^2+^ binding to EF3 and EF4 in CaM is essential for promoting the increased Po. There was no detectable effect on surface expression of Ca_V_1.2 by either WT CaM or CaM_34_ (Fig 4C,D; *Suppl*. Tab. 5). Given the ~20-fold higher expression levels of CaM_34_ versus endogenous CaM, it seems especially remarkable that this overexpression had no effect at all on Po when a lesser degree of overexpression of WT CaM induced a ~3 fold increase in Po (Fig. 4). Collectively, these data indicate that binding of Ca_2_/CaM and not apoCaM to Ca_V_1.2 at basal Ca^2+^ concentrations mediates the observed increase in Po.

**Figure 4.**
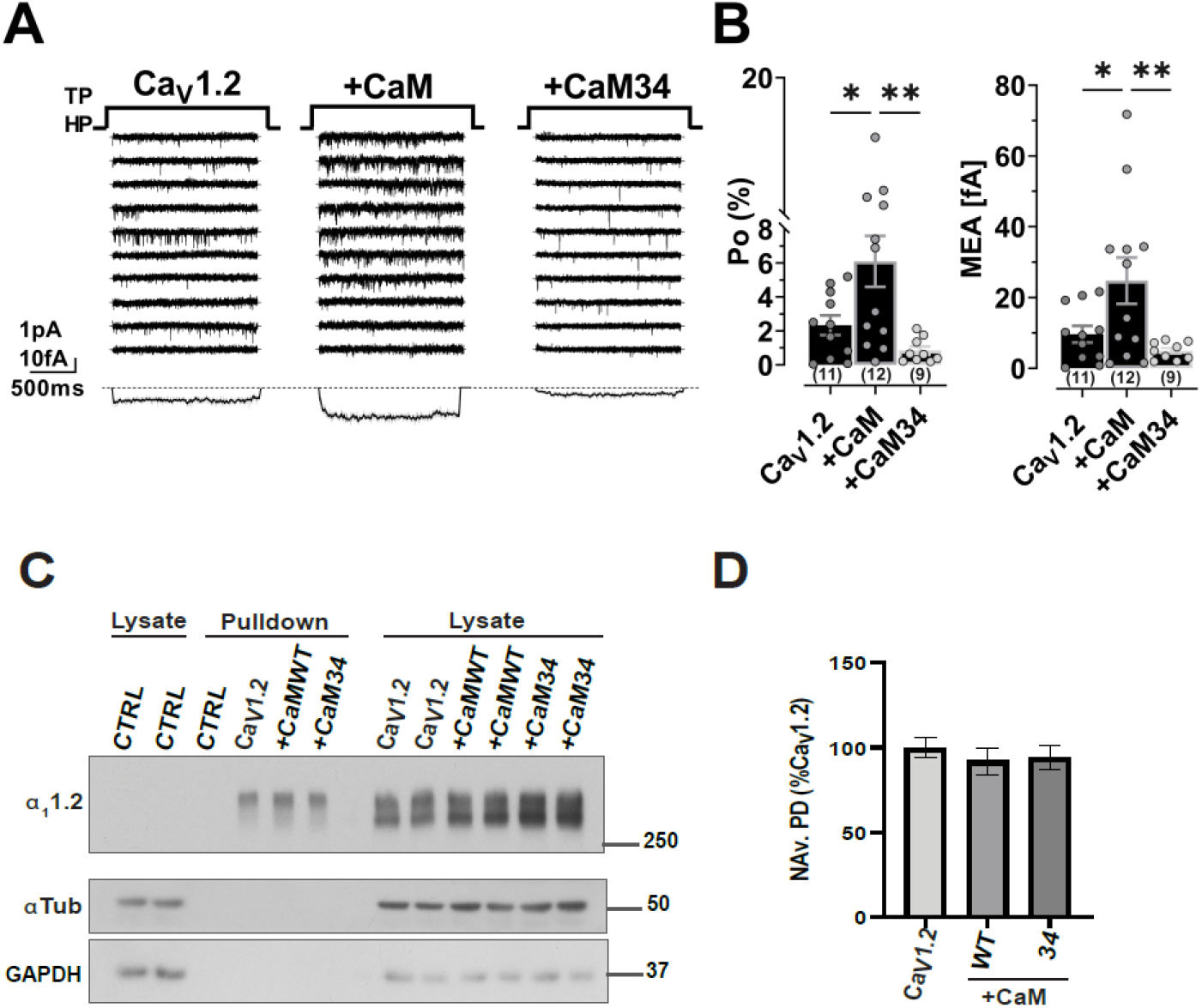
Effects of ectopic expression of WT CaM and CaM_34_ on Ca_V_1.2 activity, inactivation, and surface expression. HEK 293T/17 cells were transfected with α_1_1.2, α_2_δ1, and β_2A_ plus, if indicated, WT CaM or CaM_34_. (**A**) 10 consecutive representative single-channel traces of WT Ca_V_1.2 expressed alone (left) or together with WT CaM (middle) or CaM_34_ (right). Bottom: MEA calculated from a total of 2009 superimposed traces for Ca_V_1.2 expressed without CaM (n=11 cells), 2327 traces for Ca_V_1.2 expressed with WT CaM (n=12 cells), and 1655 traces for Ca_V_1.2 expressed with CaM_34_ (n=9 cells). (**B**) Quantification of Po (left) and MEA (right) reveal a strong increase in channel activity for ectopic expression of WT CaM but not CaM_34_ (numbers in parenthesis under bars reflect n independent recordings and error bars SEM; *p<0.05 and **p<0.01, One-Way ANOVA with Bonferroni correction). (**C**) Surface biotinylation of Ca_V_1.2 was followed by Neutravidin pulldowns (middle of blot) and immunoblotting with antibodies against the proteins indicated at the left. Right side shows respective total lysate samples in duplicate and left side total lysate samples from mock transfected cells. Cells expressing Ca_V_1.2 only or CFP-tag empty vector only were used as controls (CTRL). Tubulin (α-Tub) and GAPDH were used as loading controls for lysate samples and assessment of membrane integrity for pull down samples. Absence of tubulin and GAPDH immunoreactivity ruled out biotinylation of intracellular proteins. (D) Quantification of α_1_1.2 immunosignals in Neutravidin pulldowns (NAv.PD) normalized to the signal in Ca_V_1.2 only transfected samples (which was set to 100%; n = 7; NS = p > 0.05, one-way ANOVA, followed by Tukey’s post-hoc test).

## DISCUSSION

Pre-association of CaM with Ca_V_1.2 and the highly homologous Ca_V_1.3 under basal conditions has been suggested to both augment channel activity at low Ca^2+^ levels (Adams et al., 2014) and facilitate rapid CDI (Peterson et al., 1999, Zuhlke et al., 1999). We provide multiple lines of evidence that Ca_V_1.2 pre-associates with half-calcified Ca_2_/CaM that contains two Ca^2+^ bound to the CaM C-lobe. The fact that the CaM_34_ mutant abolished the 300% increase in channel open probability of Ca_V_1.2 caused by wild type CaM (Fig. 4B) implies that Ca^2+^ binding to EF3 and EF4 (hence half-calcified CaM) is essential for Ca_V_1.2 channel function. Also, our binding analysis reveals that IQ binding to CaM increases the apparent Ca^2+^ affinity by at least 10-fold (see *Scheme 1 and Appendix Suppl. Tab. 1*), consistent with observations from previous binding studies (Evans et al., 2011, Halling et al., 2009). Hence, the IQ-bound CaM C-lobe is more than 50% saturated with Ca^2+^ at basal Ca^2+^ concentrations when CaM is saturated with the IQ peptide (*Appendix Suppl. Fig. 1*). The concentration of free endogenous CaM inside a cell is estimated to be between 50-100 nM (Wu & Bers, 2007). As Ca_2_/CaM binds to the IQ motif with a K_D_ of 16 nM, we estimate that ~50% of Ca_V_1.2 is bound to Ca_2_/CaM under basal conditions, which would put the channel regulation by CaM in the middle of its dynamic range.

The NMR structure of Ca_2_/CaM_12’_-IQ reveals that half Ca^2+^-saturated CaM (Ca_2_/CaM) has a closed conformation (Zhang et al., 1995) in the Ca^2+^-free N-lobe and a Ca^2+^-bound open conformation (Van Petegem et al., 2005) in the C-lobe (Fig. 1). The N- and C-lobe structures of Ca_2_/CaM_12’_-IQ are separately folded and do not exhibit inter-domain contacts (Fig. 1C). The two separate lobes in Ca_2_/CaM_12’_-IQ are dynamically independent similar to apoCaM (Baber, Szabo et al., 2001, Tjandra, Kuboniwa et al., 1995, Zhang et al., 1995). The Ca^2+^-free N-lobe structure in Ca_2_/CaM_12’_-IQ does not interact with the IQ peptide, in contrast to the IQ contacts with the N-lobe observed in the crystal structure of Ca^2+^-saturated CaM (Fallon et al., 2009, Fallon et al., 2005, Van Petegem et al., 2005). The IQ peptide binds exclusively to the Ca^2+^-bound C-lobe of Ca_2_/CaM (Fig. 1D), whose structure is similar to the C-lobe of Ca_4_/CaM bound to the IQ (Fig. 1E) (Fallon et al., 2009, Fallon et al., 2005, Van Petegem et al., 2005). The IQ peptide bound to Ca_2_/CaM_12’_ is rotated 180° compared to the orientation of the IQ bound to apoCaM (Turner et al., 2020). The opposite binding orientation may explain in part why the IQ binds to Ca_2_/CaM with at least 100-fold higher affinity (Fig. 1F) compared to that of apoCaM (Evans et al., 2011, Turner et al., 2020). The contrasting binding orientation also suggests why the pre-association of Ca_V_1.2 with Ca_2_/CaM (rather than with apoCaM) predisposes Ca_V_1.2 for CDI. Since Ca_2_/CaM and Ca_4_/CaM both bind to Ca_V_1.2 with the same orientation, CaM can remain bound to Ca_V_1.2 upon Ca^2+^ influx to facilitate rapid CDI. By contrast, pre-associated apoCaM would first need to dissociate from Ca_V_1.2 upon Ca^2+^ influx and then subsequently re-bind in the conformation adopted by Ca^2+^-saturated Ca_4_/CaM to engage CDI (Fallon et al., 2009, Fallon et al., 2005, Van Petegem et al., 2005). This unbinding of apoCaM and rebinding of Ca_4_/CaM would likely prevent rapid CDI and defeat the purpose of the CaM pre-association.

Our functional analysis fully supports the relevance of prebinding of Ca_2_/CaM to the Ca_V_1.2 IQ motif. The K1662E mutation, which impaired binding of apoCaM (Turner et al., 2020) but retained binding to Ca_2_/CaM at physiological CaM concentrations of ~100 nM (Wu & Bers, 2007) (Fig. 1F), did not affect Po (Turner et al., 2020), CDI, I_Ba_ or I_Ca_ (Fig. 2). Furthermore, the Y1657D mutation impaired binding of apoCaM (K_D_ = 60 μM, Fig. 1F) as well as Ca_2_/CaM (K_D_ = 8 μM, *Appendix* Table S2), and reduced Po, CDI, I_Ba_, and I_Ca_ (Fig. 3). We also tested the effect of ectopic expression of CaM_34_ and CaM_1234_. Consistent with the earlier work on Ca_V_1.3 (Adams et al., 2014), overexpression of WT CaM strongly augmented Po (Fig. 4). However, neither CaM_34_ (Fig. 4) nor CaM_1234_ (*Appendix* Suppl. Fig. 5) increased Po despite the fact that the exogenous CaM levels were much higher (by 20-fold) than that of endogenous CaM. Accordingly, we propose that pre-association of Ca_2_/CaM and not apoCaM is important for augmenting Po and mediating CDI. Our data are consistent with earlier work on the closely related Ca_V_1.3 that showed lowering the affinity for CaM decreases its Po (Adams et al., 2014). These authors concluded that CaM in its apo form alone is what binds to the IQ motif under resting Ca^2+^ concentrations to augment Po. Their main finding was that substitution of the eponymous Ile by Met, which reduces binding of apoCaM and Ca_2_/CaM, reduced Po and overexpression of WT CaM rescued this loss. These authors provided data on the effect of membrane-tethered CaM_1234_ on whole cell currents. They did not describe the effect of untethered CaM_34_ or CaM_1234_ on Po, which would have had required cell attached single channel recording. Thus, their findings do not rule out that Po is driven by pre-association of Ca_2_/CaM. By contrast, the differential effects of: 1) the K1662E mutation on Ca_V_1.2 binding to apoCaM versus Ca_2_/CaM, 2) K1662E versus Y1657D on Po and CDI, and 3) WT CaM versus CaM_34_ or CaM_1234_ on Po collectively indicate that pre-associated Ca_2_/CaM is an important factor in determining channel Po.

As discussed above, we estimate that ~50% of Ca_V_1.2 is occupied by Ca_2_/CaM with little occupancy by apoCaM due to its low concentration in the cytosol (50-100 nM (Wu & Bers, 2007)) and low affinity binding to the IQ (K_D_ = 10 μM (Turner et al., 2020)) and full-length Ca_V_1.2 (K_D_ = 1 μM (Erickson et al., 2003)). How then can the remainder of the Ca_V_1.2 population possess a reasonable level of activity? We previously found that binding of α-actinin to the IQ motif also strongly augments Po (Turner et al., 2020). Thus, we propose a model in which Ca_V_1.2 is either occupied by α-actinin, which at the same time anchors Ca_V_1.2 at the cell surface and especially in dendritic spines where α-actinin is concentrated (Hall, Dai et al., 2013) or by Ca_2_/CaM. Accordingly, in addition to strongly promoting Po, α-actinin also augments the Ca_V_1.2 surface expression (Turner et al., 2020), perhaps by connecting to F-actin (Johnson & Byerly, 1993). On the other hand, Ca_2_/CaM augments Po with apparently little if any effect on surface expression. Channel occupancy by Ca_2_/CaM could be increased upon modest increases of basal Ca^2+^ influx potentially in a positive feedback loop at low Ca^2+^ levels and low channel activity. However, prolonged displacement of α-actinin by Ca_4_/CaM also triggers endocytosis of Ca_V_1.2 as a negative feedback mechanism (Hall et al., 2013). At this point we cannot be certain about how α-actinin and CaM intersect at the IQ motif to govern Ca_V_1.2 activity and much needs to be learned with respect to the exact function of these interactions.

In conclusion, our analysis provides novel mechanistic insight into pre-association of CaM with Ca_V_1.2 and its role in controlling channel activity and CDI. These findings are not only of functional relevance for understanding the physiological effects of Ca_V_1.2 but also inform the current understanding of pathological events such as arrhythmias due to impaired CDI (Jensen, Brohus et al., 2018, Wang, Holt et al., 2018)

## MATERIALS AND METHODS

### Cell culture and transfection of plasmids DNAs

Culture of HEK 293T/7 cells, methods for transfection and the Ca_V_1.2 expression vectors used for electrophysiological and biochemical experiments are essentially as previously described (Tseng, Henderson et al., 2017, Turner et al., 2020) and detailed in the ***SI Appendix***. Site-directed mutagenesis of the α_1_1.2 ***Y1657D*** cDNA expression vector was done as described (Tseng et al., 2017, Turner et al., 2020). The CaM expression plasmids (Peterson et al., 1999), a kind gift of J.P. Adelman, are described in the ***SI Appendix***.

### NMR spectroscopy and NMR structure calculation

All NMR measurements were performed using a Bruker Avance III 600 MHz spectrometer equipped with a four-channel interface and triple-resonance cryoprobe (TCI). NMR samples were prepared as previously described (Salveson et al., 2019). The NMR-derived structures of Ca_2_/CaM_12’_ bound to the IQ peptide were performed by restrained molecular dynamics simulations within Xplor-NIH. For further details please see ***SI Appendix***.

### Electrophysiological Recordings

Barium and Calcium currents of Ca_V_1.2 L-type Ca^2+^channels were recorded in the whole-cell and in the cell attached configuration as described previously (Turner et al., 2020). For a more detailed description of the methods and data analysis please see ***SI Appendix***.

### Surface Biotinylation and Immunoblotting

Surface biotinylation and immunoblot analysis of Ca_V_1.2 surface expression was carried out as previously described (Tseng et al., 2017, Turner et al., 2020) with the exception that biotin labeling, quenching and cell lysis were done directly on plated cells. A detailed description of this and immunoblotting methods and analysis are detailed in ***SI Appendix***.

## ACKNOWLEDGEMENTS

We thank Derrick Kaseman, Jeff Walton and Ping Yu and the UC Davis NMR Facility for help with NMR experiments. This work was supported by NIH grants R01 HL121059 (MFN), R01 EY012347 and R01 GM130925 (JBA), and R01 AG055357 and R01 NS123050 (JWH). AMC was supported by R25 GM056765 and T32 GM113770 and ZME-T by T32 GM 007377.

## AUTHOR CONTRIBUTIONS

M.C.H, J.W.H. and J.B.A. designed research and, with input from other authors, wrote the paper; P.B., I.S., A.M.C., D.E.A., G.J., Z.M.E.-T., K.N.M.M., M.N.-C., M.F.N., M.C.H. and J.B.A. performed experiments; P.B., I.S., A.M.C., D.E.A., Q.Y., E.K., M.N.-C., M.F.N., M.C.H, J.W.H., and J.B.A. analyzed data.

## DATA AVAILABILITY

Atomic coordinates were deposited in the Protein Databank (accession no. 7L8V).

